# Gene Expression Signatures of Sporadic ALS Motor Neuron Populations

**DOI:** 10.1101/038448

**Authors:** Ranjan Batra, Kasey Hutt, Anthony Vu, Stuart J. Rabin, Michael W. Baughn, Ryan T. Libby, Shawn Hoon, John Ravits, Gene W. Yeo

## Abstract

**Background:** Amyotrophic lateral sclerosis (ALS) is a fatal neurodegenerative disease primarily affecting motor neurons (MNs) to cause progressive paralysis. Ninety percent of cases are sporadic (sALS) and ten percent are familial (fALS). The molecular mechanisms underlying neurodegeneration remain elusive and there is a lack of promising biomarkers that define ALS phenotypes and progression. To date, most expression studies have focused on either complex whole tissues that contain cells other than MNs or induced pluripotent derived MNs (iMNs). Furthermore, as human tissue samples have high variability, estimation of differential gene-expression is not a trivial task.

**Results:** Here, we report a battery of orthogonal computational analyses to discover geneexpression defects in laser capture microdissected and enriched MN RNA pools from sALS patient spinal cords in regions destined for but not yet advanced in neurodegenerative stage. We used total RNA-sequencing (RNA-seq), applied multiple percentile rank (MPR) analysis to analyze MN-specific gene-expression signatures, and used high-throughput qPCR to validate RNA-seq results. Furthermore, we used a systems-level approach that identified molecular networks perturbed in sALS MNs. Weighted gene co-expression correlation network (WGCNA) analysis revealed defects in neurotransmitter biosynthesis and RNA-processing pathways while gene-gene interaction analysis showed abnormalities in networks that pertained to cell-adhesion, immune response and wound healing.

**Conclusions:** We discover gene-expression signatures that distinguish sALS from control MNs and our findings illuminate possible mechanisms of cellular toxicity. Our systematic and comprehensive analysis serves as a framework to reveal expression signatures and disrupted pathways that will be useful for future mechanistic studies and biomarker based therapeutic research.

## Introduction

Amyotrophic lateral sclerosis (ALS) is a progressive fatal neuromuscular disease that predominantly affects upper and lower motor neurons [1] [2]. The majority (90%) of cases are sporadic (sALS), and a minority (10%) of cases are familial (fALS) [3]. The causes of sporadic ALS and the mechanisms of neurodegeneration even in genetically defined cases are unknown. The most common neuropathological hallmark of sALS is TDP-43 positive cytoplasmic inclusions, especially in neurons [4]. At present, there is no cure for ALS and a dearth of clinical biomarkers for the development of effective clinical targets. Studying geneexpression and RNA-processing in the patient tissues can lead to identification of transcriptional and post-transcriptional disturbances and relevant biomarkers, and even novel mutations that define the strikingly similar phenotypes of this diverse group of patients. Importantly, there appears to be a central role for the disruption of RNA processing in ALS with the discovery of fALS mutations in RNA binding proteins (RBPs) including *TARDBP* (TDP-43), *FUS, TAF15, HNRNPA2B1, HNRNPA1, EWSR1* and *ATXN2*. Thus the strategy to identify RNA signatures delineating ALS seems a promising one.

Recently, there has been a number of studies that have profiled human tissues [5] [6] [7] [8]. These studies profiled gene-expression signature in homogenized whole tissues [9]. Although these approaches are useful for discovering mechanisms of pathogenesis and druggable targets, they overly simplify the reality of complex heterogeneous tissues such as the nervous system. One recent study was carried out for *C9orf72* ALS (C9ALS) and sALS in frontal cortex and cerebellum and identified many ALS-related expression and splicing changes [9]. Surprisingly, cerebellum, a region of the central nervous system (CNS) thought relatively spared clinically in patients, was particularly affected in C9ALS patients. However, the study does not assess transcriptome-wide changes in a particularly disease-relevant cell population, such as motor neurons in patient spinal cords. A number of profiling studies have selectively enriched RNA pools, such as dissecting the anterior horns [10] or by microdissection [11] [12] [13] [14]. In our previous study, we used exon oligonucleotide microarrays to profile gene-expression in sALS motor neurons (MNs) using laser capture microdissection (LCM) [12]. Another study analyzed a smaller number of sALS samples using Affymetrix exon arrays to profile LCM lower motor neurons [14]. These studies reported alterations in splicing and gene-expression profiles in ALS but a comprehensive systems-based gene-expression analysis was only carried out for *C9orf72* ALS [15]. Furthermore, it is difficult to establish if the gene-expression changes are the result of neurodegeneration.

Clinical observations in ALS suggest that neurodegeneration may begin focally and progress along anatomically defined neuronal tracts over the disease path [16, 17]. The site of onset in ALS is often predictive of the neuroanatomical path of degeneration [16]. This creates an opportunity for predicting and isolating neurons that lie along the course of future neurodegeneration but that have not succumbed to disease at the time of death when the disease process subsumed respiratory motor neurons anatomically proximal in the disease path [17]. Here we use RNA-sequencing methodology to define transcriptomic signatures of sALS spinal motor neurons (not succumbed at the time of isolation) isolated by laser capture microdissection, extending our previous microarray study [12]. We use three main statistical and analytical approaches to report unique transcriptomic signatures of these neurons that are different from those described by whole tissue analysis of different brain regions, or LCM captured MNs from C9orf72 ALS patients. Network based analysis of expression data identified specific modules of genes that are affected in the sALS neurons including immune response and cell adhesion. In addition, gene interaction network analysis identified defects in RNA processing. Our data and findings will serve as an important framework for uncovering mechanisms underlying sALS, identifying candidate targets for therapy, and discovering candidate biomarkers.

## Results

### Laser capture microdissection results in high quality collection of RNA

To obtain transcriptome-wide gene expression measurements in adult spinal motor neurons (MNs), we performed laser capture microdissection (LCM) of lumbar spinal cord sections from 13 sALS patients and 9 control patients (Figure 1A, Supplementary Table 1A). Nervous systems selected for profiling for sALS were from patients who had bulbar or arm onset of disease and caudally progressing disease and thus had abundant residual motor neurons in the lumbar region at the time of death. Total RNA was extracted and amplified using random priming, converted into cDNA, linearly amplified, fragmented and subjected to sequencing library preparation. The quality and quantity of products was controlled at each step using microelectrophoresis and RNA integrity numbers (Supplementary Table 1) and resulted in high-quality libraries that were submitted for high-throughput Illumina sequencing.

**Figure 1.**
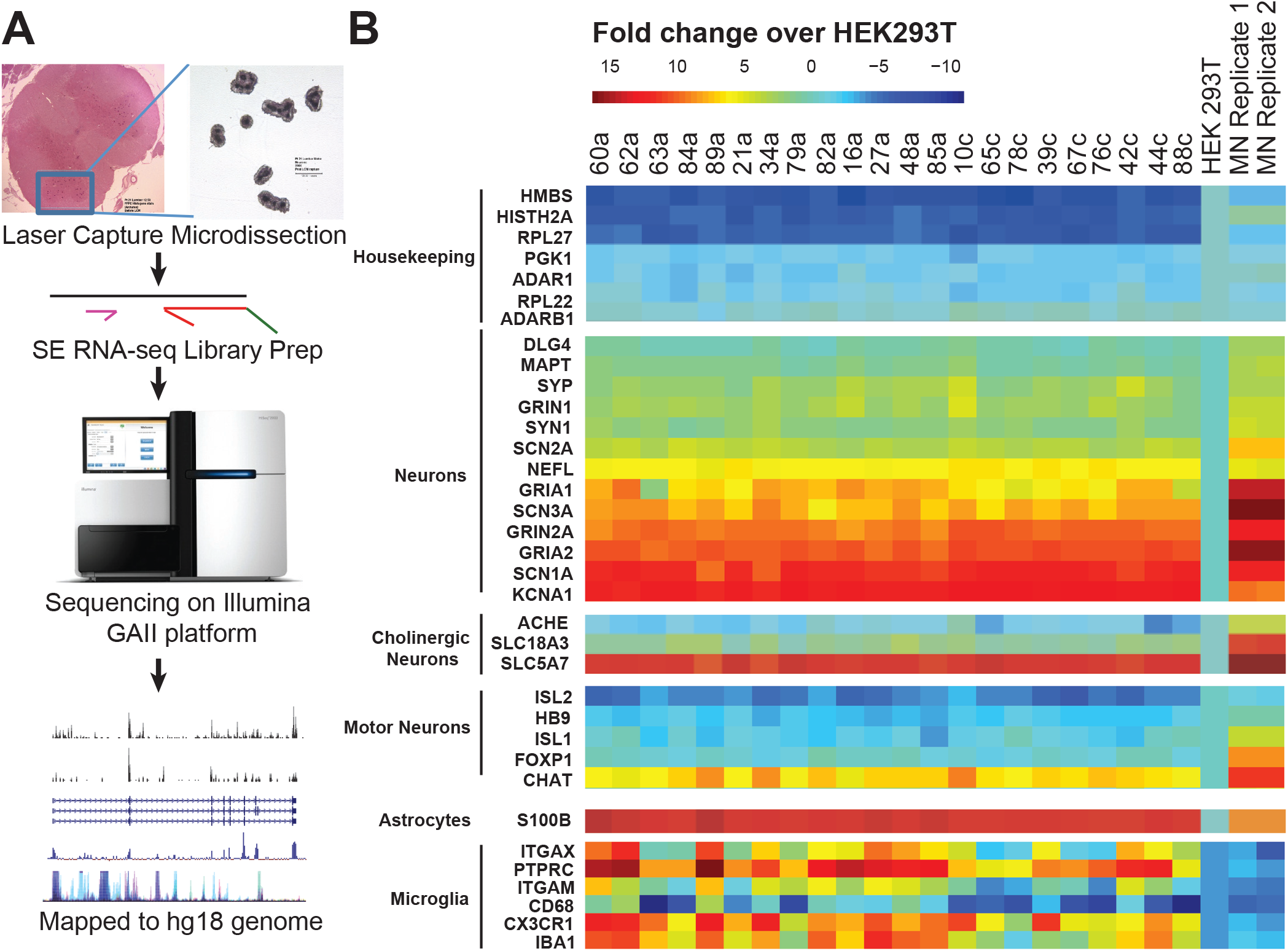
RNA-seq data analysis shows enrichment for motor neuron (MN) signature in laser capture microdissected motor neurons from sporadic ALS and control spinal cords. (A) Strategy for differential gene expression analysis of LCM isolated MNs. (B) Heatmaps of relative expression levels of housekeeping genes (top), neuronal makers (middle), and glial markers (bottom) in MNs from spinal cords compared to human 293T cells and in vitro derived MNs from normal iPSCs (last two lanes).

### RNA-seq quality assessment shows sufficient sequencing depth and specific enrichment for motor neuron signature

The 22 RNA-seq libraries were randomized (Supplementary Table 1B) and sequenced on Illumina GA II to an average depth of ∽28 million reads per sample (Table 1). Reads were mapped to the hg18 genome assembly using the alignment software Bowtie [18] and subsequent gene assignment demonstrated an approximately four-fold enrichment for reads overlapping annotated gene boundaries over intergenic space, indicating that RNA extraction was sufficient in capturing ∽5000 (log2RPKM > 1) annotated protein-coding genes. To evaluate if the capture enriched for motor neurons (MNs), we included RNA-seq data from motor neurons derived from two independent induced pluripotent stem cells (iPSCs) as comparison for an early MN signature (Figure 1B, right two lanes), as well as human epithelial kidney cells (HEK293T) as a non-neuronal comparison. Essentially all the genes within our neuronal panel were enriched in the LCM-isolated MN samples over the HEK293T cells. This included the N-methyld-aspartate (NMDA) receptor *GRIN2A* and the alpha-amino-3-hydroxy-5-methyl-4-isoxazole propionate [4] receptor *GRIA2* (Figure 2B). The marker genes typical of motor neurons, *CHT1, CHAT*, and to a lesser extent, *NKX6-2* were enriched (Figure 1B and Supplementary Figure 1). Early MN markers *ISL1* and *ISL2* were not overrepresented, indicating that the LCM-isolated MNs harvested were mature. Somewhat surprisingly, *HB9* was also not enriched when compared to 293T cells, although our data suggests this gene is also expressed in 293T cells. An astrocyte marker *SB100* and microglial markers like *PTPRC* and *CX3CR1* were also enriched in our MN pool (Figure 1B). We concluded that our LCM followed by RNA-seq approach was successful in obtaining a MN-enriched population of cells from sALS and control patients, recognizing that other cell types such as astrocyte and microglia RNA were also unavoidably present.

**Figure 2.**
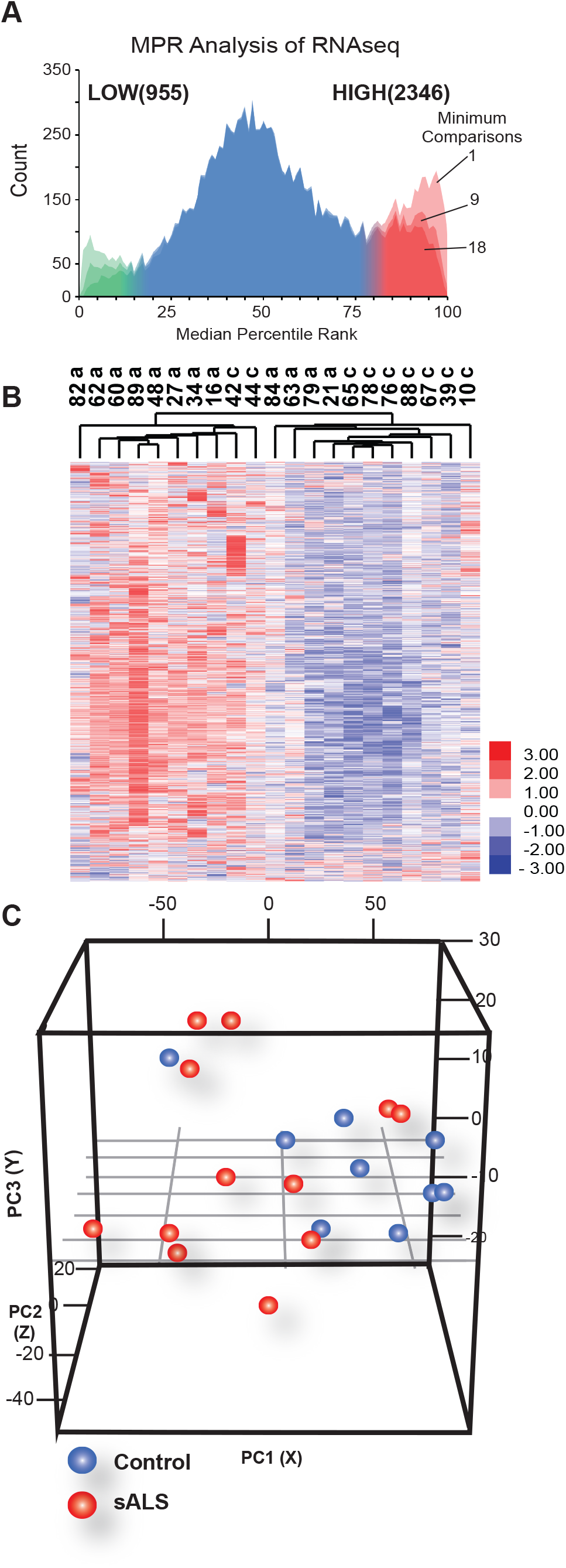
Gene expression analysis reveals differential signature in sporadic ALS samples. (A) Median Percentile Rank (MPR) analysis reveals genes that are relatively higher (red, n =2346) and lower (green, n=955) expressed in sALS versus control. (B) Hierarchical clustering of gene RPKMs of differentially expressed genes across patient samples (sALS, N=12, Control, N=9). (C) Principal component analysis (PCA) of differentially expressed genes across patient samples (sALS, N=12, Control, N=9).

### RNA-seq data analysis of sALS motor neurons identifies unique gene-expression signature

To identify expression differences in these highly enriched MN samples that distinguish sALS patients from controls, we applied a modification of a published method [19], termed <u>M</u>edian <u>P</u>ercentile <u>R</u>ank (MPR). More frequently used methods that identify differentially expressed genes, such as EdgeR [20] and DE-seq [21], are sensitive to the naturally higher sample variability from human samples. Therefore, we utilized MPR which was previously used to find enriched targets in comparably noisy data generated by ChIP-on-chip [19] (see methods). Briefly, the MPR approach begins by ordering the genes by its fold-change in each pairwise comparison between sALS and control samples. Next, the percentile rank of each gene is calculated based on the position of the gene in the ordered (high to low fold-change) list. The MPR value is then calculated as the median of percentile ranks for a given gene across all comparisons. The MPR values for all genes is displayed using a histogram, and any gene that is significantly upregulated or down-regulated is positioned towards the edges of the histogram, and cutoffs are estimated empirically from the proportion of genes occurring near the edges. From the 117 pairwise comparisons, we observed that two clear cutoffs appear (Figure 2A and Supplementary Table 2A, cutoffs of 0.85 and 0.15). These cutoffs produced 2,346 upregulated and 955 down-regulated genes in sALS patient compared to control samples. Interestingly, the upregulated set of genes (in red) was more robust to increasingly conservative (e.g. requiring that the gene is counted only in higher numbers of comparisons) iterations for MPR inclusion (Figure 3A, darker shades of red or green) than the down-regulated set, suggesting that the upregulated genes are more robustly affected between comparisons of sALS samples to control samples. Hierarchical clustering of the expression values of the 1500 genes (RPKM values higher than 2.0 and max. - min. >2.0, Supplementary Table 2D) that were predicted to be distinct between sALS and controls by MPR analysis led to the clear segregation of at least 8 out of 12 sALS and at least 7 out of 9 control samples (Figure 2B). Furthermore, principal component analysis (PCA) of these samples based on MPR-identified genes confirmed that sALS samples were indeed much more variable than the control samples (Figure 2C). To evaluate the molecular and biological processes that are altered, we performed Gene Ontology analyses using the DAVID package [22]. We identify enrichment in extracellular region (P value = 5.3E-31), immune response (*P* value = 5.1E-27), and biological (cell) adhesion (P value = 7.7E-21) categories (Supplementary Table 2) in the genes upregulated in sALS samples. In summary, our results demonstrated that sALS motor neuron enriched RNA pools have a differential RNA signature compared to control patients.

**Figure 3.**
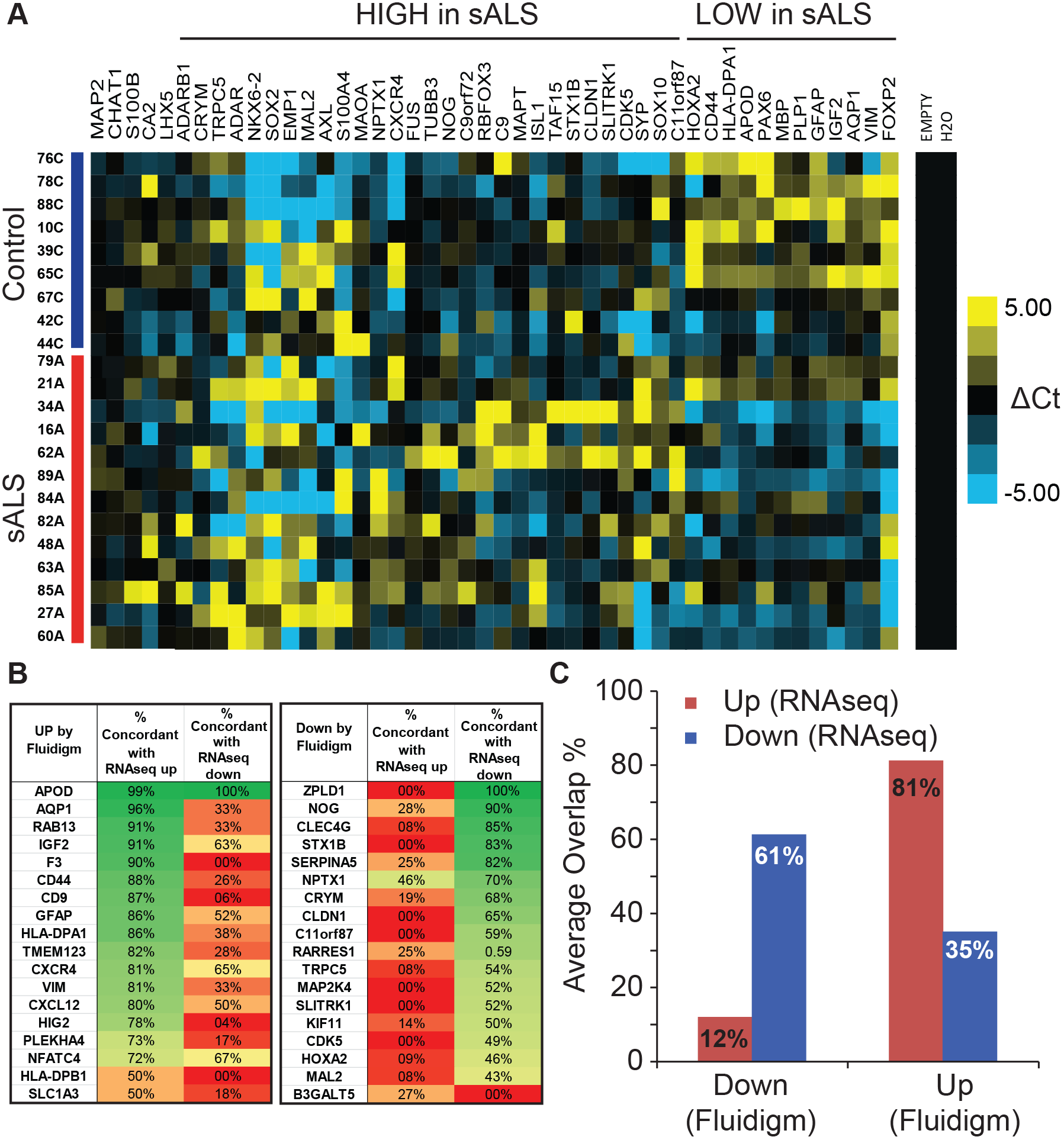
High-throughput validation of RNAseq data using Fluidigm Biomark qPCR system. (A) Normalized heatmap of Ct values for qPCR primer pairs on Fluidigm Biomark qPCR platform. The dCt values were estimated by normalizing to 4 uniformly expressed house keeping genes *(ACTB, RPL27, GAPDH*, and *PGK1)* (B) Overlap statistics between qPCR pairwise fold-changes and RNAseq pairwise fold changes. Two charts show percentage of genes that overlap. Concordant overlaps (up-up and down-down) are green, discordant overlaps (up-down, down-up) are red. (C) Summary of overlap charts reveals upregulated genes by RNA-seq (red bar) show much higher validation rate than down-regulated genes by RNA-seq (blue).

### Gene-expression validation using high-throughput qPCR shows high congruence

As an independent method to validate the expression changes discovered by RNA-seq, we performed quantitative RT-PCR using the Fluidigm Biomark platform. The 22 original cDNA samples were interrogated with a selection of 18 upregulated and 18 down-regulated genes, as well as a smaller panel of genes predicted to be unchanged (control genes). A heat map of delta Ct values (Ct value of target gene - Ct value of control genes) is shown (Figure 3A). Our results demonstrate that our panel clearly distinguishes between sALS and control cells, while also visibly reflecting the natural variation in these human samples (Figure 3A). To evaluate how well qRT-PCR compares with our RNA-seq and MPR-based predictions, each qRT-PCR pairwise comparison was validated against the same RNA-seq paired comparison. For each pair of sALS and control sample compared, the qRT-PCR derived fold-change of the gene was compared with its fold-change as measured by RNA-seq. If the expression of the gene was conservatively two-fold different between sALS and control sample by both qPCR and RNA-seq, we first considered that the methods reliably detected the change in the target gene. Next, if the gene is changing in sALS relative to control sample in the same direction by both qPCR and RNA-seq, the gene would be considered concordantly validated and is highlighted in green in Figure 3B. But, if the qRT-PCR comparison yields discordant directions of change by RNA-seq, it would be highlighted in red (Figure 3B). The percentage of reliably detected changes between all pairwise comparisons where the direction of change (i.e. higher in sALS) was the same was determined for each gene evaluated (Figure 3B). Notably, the down-regulated genes showed lower actual agreements, as the most highly predicted down-regulated gene was concordant in only 52 pairwise comparisons (Figure 3C). In contrast, the most upregulated gene was concordant in 87 pairwise comparisons, which suggests that the upregulated set has higher predictive value in distinguishing disease versus control expression changes (Figure 3C). In summary, we conclude that at least ∽80% and ∽60% of the upregulated and downregulated genes, respectively, are concordant between both technologies.

### Systems-level analysis of sALS motor neuron gene-expression reveals affected cellular networks

To assess what genes in our MPR dataset interacted with each other based on experimental evidence of co-expression, neighborhood and pathways, we used the String v10 software to assemble them into high confidence (0.90) biological networks (Figure 4A, full network in Supplementary Figure 2, network coordinates in Supplementary Table 3E) [23]. We identified 4 distinct groups of high confidence interaction networks, which were empirically annotated as (1) cell adhesion and extracellular matrix, (2) chemokines and immune response, (3) cell signaling, and (4) glutamate and neurotransmitter metabolism. We found that intersection of sALS MN gene-expression with sALS and C9ALS cerebellum and frontal cortex data [9] showed little overlap implying that different regions of the adult CNS may be distinctly affected and mixed populations of cells are likely to reveal different gene-expression signatures (Supplementary Figure 3).

**Figure 4.**
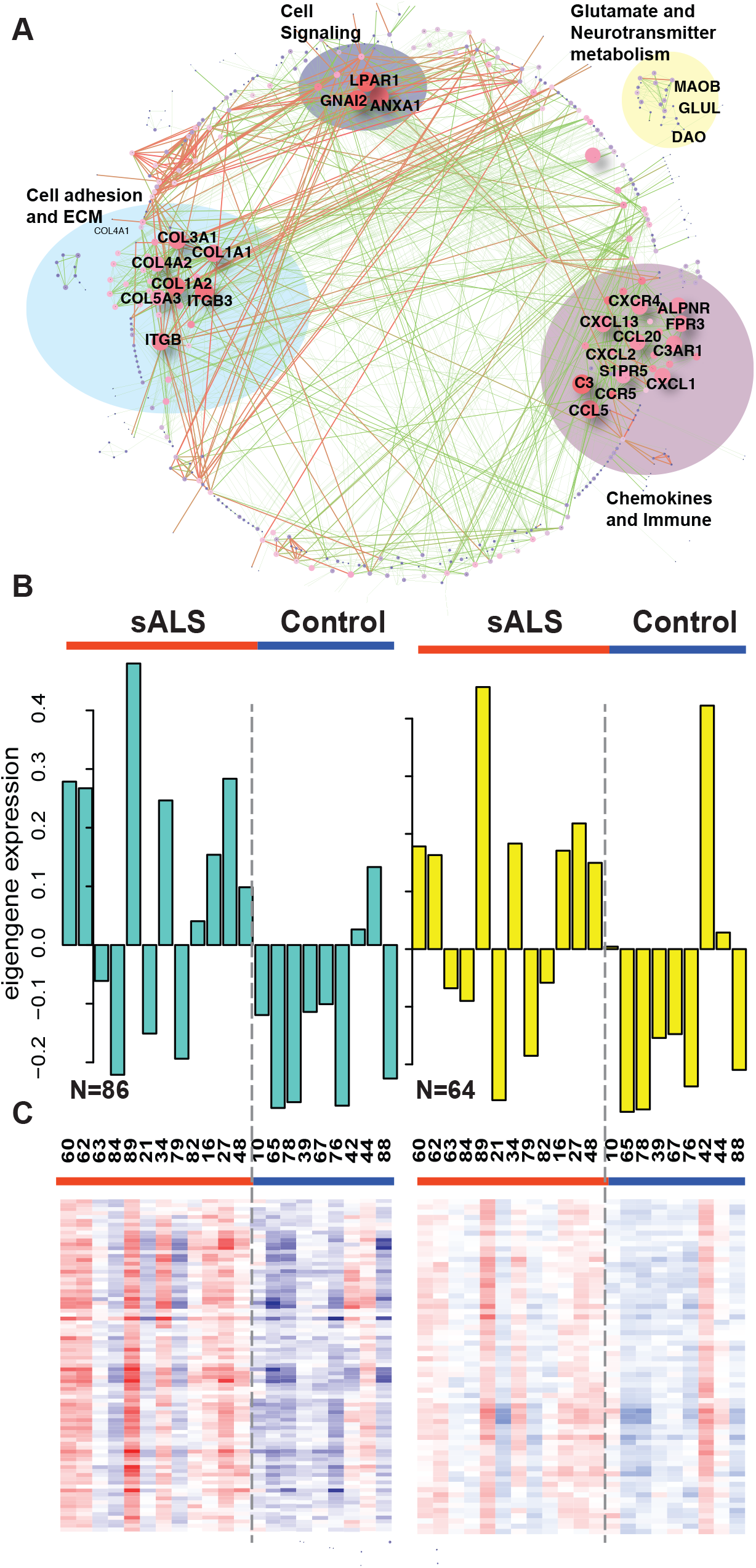
Systems analysis reveal modules of aberrantly regulated genes in sporadic ALS. (A) Gene interaction network analysis using STRING v10 tool shows many overlapping pathways. The nodes are colored according to degree or number of interactions for each node (red = high; dark blue = low). Red nodes are therefore hubs and are predicted to effect downstream pathways more. Edges are colored according to experimental co-expression (red = high; green = low). Red edges connect genes that are more likely to be co-expressed or co-affected. Prominently affected pathways are shown. (B) Bar charts showing representative eigengene values (first principal component and representative value for expression of the whole module) of the turquoise and yellow modules upregulated in sALS. (C) Heatmap of RPKM values of genes in turquoise and yellow modules.

Another avenue to gain insights into the pathways that are distinct between the expression profiles of disease and control conditions is by employing weighted gene co-expression correlation network analyses (WGCNA) [24] [25]. This analysis focuses on a module or a group of genes (rather than a single gene) that appear to be correlated in their expression. The genes are hierarchically clustered and the branches of the dendrograms with similar geneexpression are designated as modules. We applied the WGCNA co-expression analysis to our LCM MN RNA-seq data. The MN gene dendrogram had only a few distinct branches and 3 significant modules altered in sALS samples were detected (Figure 4B and Supplementary Figure 4, Supplementary Table 3A). Modules (named as colors, for ease, inherently by the software) were represented by their module eigengene (ME) values, which is the first principal component that represents the highest percent of variance for all genes in a module. One module contained only ribosomal RNA and related proteins and it could be attributed to either mild ribosomal RNA contamination or increased translation. We assumed rRNA contamination as total RNA and random priming were used for library generation and discounted this module. The two other significant modules were annotated using DAVID (Supplementary Tables 3C and 3D). The turquoise module (N=86) was enriched for GO categories organic acid biosynthetic process, carboxylic acid biosynthetic process (includes neurotransmitter biosynthesis), and positive regulation of cell communication The yellow module (N=64) was enriched for the GO categories RNA splicing, mRNA metabolic process, and mRNA processing. Interestingly this RNA processing module includes genes such *HNRNPA2B1*, and *MALAT1*, which have previously been implicated in ALS (Supplementary Figure 4) [26], [27], [28]. Other RNA binding proteins (RBPs) such as hnRNPC, SFRS12, RBM5 are also present in the RNA processing yellow module. Genes expressed in both these modules were higher in the sALS relative to control samples (Figure 4B and 4C), in agreement with our MPR analysis that the upregulated genes were more robust in separating sALS from controls. Interestingly, only 13 genes were shared between the genes found upregulated by MPR (2,346 genes) and the modules identified by WGCNA, consisting of 150 genes.

## Discussion

In this study, we examined gene-expression differences in LCM enriched MN populations that were destined for degeneration to discover RNA signatures that underlie ALS molecular pathogenesis. This approach enriches for molecular changes that are relatively upstream rather than late in the neurodegeneration cascade, although it is probable that the molecular cascade leading to cell death had already begun. Our approach to enrich for MN populations has important advantages over profiling a degenerated and homogenized complex tissue sample with different cell populations. Our MN populations were enriched for mature neuronal markers and subsequent analysis of differential expression revealed 2,346 upregulated and 955 down-regulated genes. We used the MPR method for analyzing differential expression instead of more popular methods such as EdgeR and DE-seq. While conventional methods use either Poisson or negative binomial distributions followed by estimation of differential expression using the Benjamini-Hochberg corrections [29], [21], MPR involves multiple pair-wise comparisons (9 controls x 13 sALS = 117 comparisons) to calculate the median percentile ranks for each gene. Based on independent validation using the Fluidigm platform, we found that the MPR method worked reasonably well to identify robust expression changes when the data variability is high, such as in studies involving collected human tissue samples.

Our RNA signature of ∽1500 genes segregated the sALS and control MN groups confirming that the ALS condition is the driving factor that resulted in differential signatures between sALS and control MNs. High-throughput qPCR based validation allowed us to validate 96 gene pairs in a single run and our qPCR data was highly consistent with our RNA-seq based detection of gene expression changes (Concordance = 80% and 60% for upregulated and downregulated genes respectively). Interestingly, ALS related *C9orf72* and *FUS* genes were slightly upregulated in sALS samples (Figure 3A). GGGGCC repeat expansions in the first intron of the *C9orf72* gene are known to be the most common cause of fALS [30] [31] and mechanisms including RNA-processing defects, RAN protein toxicity, and *C9orf72* haploinsufficiency have been implicated for *C9orf72* ALS [32] [33] [34] [3]. Mutations in RBP encoding gene fused in sarcoma (FUS) are also associated with fALS and changes in FUS levels are thought to affect RNA-processing in ALS [35] [36] [37]. Another RBP gene *RBFOX3*, whose protein product is recognized by the neuronal marker NeuN antigen, was also found to be upregulated in sALS MNs. Curiously, microtubule associated protein tau *(MAPT)* also appeared to be upregulated in some sALS cases (Figure 3A). This is intriguing as Tau is associated with Alzheimer’s disease and mutations in *MAPT* cause frontotemporal dementia with parkinsonism 17 (FTDP-17) [38] [39]. Noteworthy among the genes downregulated in sALS were apolipoprotein D (APOD), which is associated with aging [40]; paired box 6 (PAX6), important neuronal development; and forkhead box P2 (FOXP2), important for speech and language, mutations in which are associated with speech-language disorder 1 (SPCH1) [41] [42].

Since genes function in biological networks, several systems-level computational analyses were also performed on our RNA signature to identify pathways misregulated in sALS MNs. First, we constructed co-expression matrices using WGCNA and confirmed that the differentially expressed modules are upregulated in sALS. The recognized modules were robustly overexpressed in most sALS samples, in agreement with MPR-based predictions. GO annotations for upregulated modules showed enrichment for organic acid biosynthetic process (involves neurotransmitter biosynthesis) and RNA splicing/processing. Our results confirm previous reports that have associated RNA processing defects in ALS [43]. We also found previously implicated *MALAT1*, a long non-coding RNA, and *HNRNPA2B1*, an RBP gene, mutations in which are associated with familial ALS. It is also prominent that *HNRNPA2B1* is upregulated in sALS MNs. Since it also contains a prion-like domain similar to TDP-43, we plan to test the hypothesis in future that it may be involved in the process of inclusion formation in ALS. Three more ALS-associated genes were also identified by MPR list but not in WGCNA. Damino-acid oxidase gene DAO [44] and *SIGMAR1* [45] were upregulated whereas recently discovered TUBA4A [46] was downregulated. Curiously, another long non-coding RNA *HOTAIR* was downregulated that could lead to upregulation of several epigenetically silenced genes. The known targets for HOTAIR are *HOXD (HOXD10, HOXD11, HOXD3)* genes, which did not satisfy our thresholds but seemed to be moderately upregulated in our RNA-seq data [47], whereas HOXA2 was significantly downregulated as validated by qPCR (Figure 3A). MPR does not take co-expression into account whereas WGCNA is sensitive to extreme data variability, which may account for the low overlap between the two datasets. This emphasizes the importance of rigorous statistical methods, multiple approaches and extensive independent validation in analyzing gene-expression signatures in human samples. We also analyzed gene interactions on a systems level using the STRING software package [23]. Genes that interact on the basis of experimental evidence and co-expression are associated with each other within networks. We detected that genes involved in cell adhesion, extracellular matrix (ECM), and immune response were prominent which was concordant with GO analysis of MPR based differentially expressed genes [22]. Intriguingly, cell adhesion and ECM categories were enriched in MN alternative splicing (AS) data as was previously reported [12].

## Conclusions

We have uncovered gene-expression changes that occur in MNs destined for neurodegeneration in sporadic ALS. Our methods demonstrate the requirement for multiple statistical approaches and rigorous validation of human gene-expression data to successfully identify disease related transcriptome signatures. Our data not only provides potential mechanisms of cell adhesion and RNA processing that may lead cell death but also provides candidate biomarkers that are validated by an independent assay for drug screening studies in the future.

## METHODS

### Tissue acquisition and pathological screening

All nervous systems were acquired by way of an Investigational Review Board and Health Insurance Portability and Accountability Act compliant process. The sALS nervous systems were from patients who had been followed during the clinical course of their illness and met El Escorial criteria for definite ALS. Nervous systems selected for profiling for sALS were from patients who had bulbar or arm onset of disease and caudally progressing disease and thus had abundant residual motor neurons in the lumbar region at the time of death. Control nervous systems were from patients from the hospital’s critical care unit when life support was withdrawn. Upon death, autopsies were performed immediately by an on-call tissue acquisition team. Tissue collections were completed within 6 hours, usually within 4 hours, of death and the entire motor system was dissected and archived for downstream applications by creating two parallel tissue sets from alternating adjacent regions. For molecular studies, segments were embedded in cutting media, frozen on blocks of dry ice and stored at −70 C. For structural studies, the adjacent segments were fixed in 70% neutral buffered formalin, embedded in paraffin (formalin-fixed paraffin-embedded or FFPE) and stored at room temperature.

### Laser capture microdissection (LCM) and RNA extraction

Laser capture microdissection was done according to the protocol described previously (Rabin et al). Briefly, Thirty-five to 50 sections of frozen OCT-embedded human lumbar spinal cord were cut at a thickness of 9 μm in a −18C cryotome and placed onto uncharged glass slides. The sections were returned immediately to −78C after production, and maintained at that temperature for a minimum of three hours. Staining with cresyl violet acetate was accomplished in a 10-step, timed, nuclease-free immersion process. Motor neurons were microdissected from each slide using a Pixcell IIe Laser Capture Microdissection (LCM) System (Arcturus Bioscience) and CapSureTM Macro LCM Caps (Applied Biosystems). Each LCM session had an upper time limit of 2.5 hours. In addition to the motor neuron enriched LCM material, the remaining anterior horn region was collected to create a second parallel pool. RNA isolation was carried out in each using an RNeasy Micro kit (Qiagen), which employs guanadinium isothiocyanate and 2-mercaptoethanol extraction and column purification, including an on-column DNase I digestion step. Aliquots of each RNA pool were analyzed for quality and quantity on a Bioanalyzer RNA Pico chip (Agilent) and only those samples with RNA Integrity Numbers (RINs) > 5 or evidence of 28S peaks on the electropherogram tracings were advanced to the next step.

### RNA-seq library preparation and sequencing

Total RNA (10 ng) was amplified to cDNA using random priming with the Ovation^^®^^ RNA-Seq System (NuGEN). This procedure involved initial generation of double-stranded cDNA followed by amplification through production of single-stranded cDNA using the SPIA^^®^^ process. The single-stranded cDNA was then copied into double-stranded cDNA and quantified using the PicoGreen kit (Invitrogen) assayed with the FUSION system (Packard Biosciences), resulting in a total yield of 3-4 ug amplified cDNA’s. Standard concentration curves using bacteriophage lambda DNA were generated for each PicoGreen analysis, and the samples were diluted by 10fold serial dilutions. QC of the amplified cDNA was determined using a Bioanalyzer running an RNA 6000 Nano LabChip (Agilent). Double-stranded cDNA (1-2 ug) was fragmented to 150-200 bp sizes using Adaptive Focused Acoustics™ (Covaris, Inc., Woburn, MA), and QC of the fragmented cDNA was performed using a Bioanalyzer DNA Chip 1000 (Agilent). Fragmented cDNA’s were concentrated using the QIAquick PCR Purification Kit (Qiagen) and 200 ng of the fragmented cDNA’s (determined following PicoGreen quantitation) were end-repaired, followed by adaptor ligation onto the fragments and amplification using the Encore^^®^^ NGS Library System I (NuGEN). Library QC was performed using a Bioanalyzer DNA Chip 1000.

### Gene-expression analysis of RNA-seq data

Raw reads were adaptor-trimmed and aligned to the human genome (UCSC version hg18) using bowtie (parameters: -l 20 -m 5 -k 5 --best -q hg18). Aligned reads were assigned to a custom annotation of human genes [36], [48], and gene expression values were calculated as RPKMs [28]. Control versus sALS patient datasets were compared in a pairwise fashion, calculating ratio fold-changes for all possible 117 disease versus control pairs. For Median Percentile Rank (MPR) analysis, each set of fold-changes were sorted and given a percentile rank, and the median percentile rank for each gene was determined and plotted as a histogram. The local minima near the down-regulated edge, 0.15 was taken as the down-regulated cutoff, and a mirrored cutoff of 0.85 was taken for the upregulated cutoff, and these gene sets were used for subsequent GO analysis. To demonstrate the validity of this analysis, significantly changed genes in each of the 117 comparisons were calculated from local Z scores as before [28], and the union of changed genes were used for hierarchical clustering (Matlab). The ordered list of genes was used to generate a sliding window calculation of MPR values and MPR-determined up- and down-regulated genes.

### Fluidigm Validation

High-throughput qPCR validation was carried out on Fluidigm’s Biomark HD system, as described in manufacturer’s protocol. Specifically, cDNA (6.25 ng) was pre-amplified using 50 nM pooled primer mixture and TaqMan PreAmp Master Mix (Applied Biosystems) for 13 cycles. Unincorporated primers were removed with 8U of Exonuclease I (NEB) and the reaction products were diluted 10-fold in TE Buffer (TEKnova). The treated cDNA’s (280 pg) were combined with SsoFast EvaGreen Supermix with Low ROX (Bio-Rad) and loaded onto the 96.96 Dynamic Array integrated fluidic circuit (Fluidigm). Primers used are provided in Supplementary table 4. The overlap of Fluidigm results with RNA-seq was performed by comparing the fold-changes found for the specific pairwise comparison. For a particular gene in a single pairwise comparison, if the RNAseq and Fluidigm fold-changes were both greater than 2-fold in the same direction, the gene was considered to be validated positive.

### Co-expression analysis using WGCNA

RPKM values of genes (log_2_RPKM > 1) were used to construct signed co-expression networks using the WGCNA package in R. Low expression genes were excluded from the analysis to remove noise to satisfy the scale free toplogy and R^2^ value of > 0.8. Networks were constructed by obtaining a dissimilarity matrix based on the topological overlap. The adjacency matrix was calculated by raising the correlation matrix to a soft power of 40 (signed network),which was chosen to attain scale-free topology. Modules were calculated by hybrid treecutting function with deepsplit parameter = 1-3. Bar graphs for representative eigengene (1^st^ principal component) values were obtained using the barplot function. Module membership (kME) was calculated as the correlation between gene-expression values and the module eigengene. Modules were visualized using Cytoscape network viewer 3.1.0, and annotated for gene ontology using DAVID. Heat maps for modules were generated using RPKM values for genes represented in the module, using Cluster 3.0 and visualized using TreeView.

### Gene-interaction network

Gene or protein-protein interaction network was constructed using String v10. Gene names were inputted and medium-high stringency was chosen for network detection based on existing co-expression, gene interaction, and experimental evidence data. Networks were visualized and colored according to degree of the nodes (node color; red:blue = high:low) and coexpression for edges (red:green = high:low) using Cytoscape 3.1.0.

## Author Contributions

J.R. and G.W.Y. conceived the study. M.W.B. and R.T.L. generated the total RNA pools. S.J.R. generated the libraries. A.V. generated the motor neurons from iPSC. S.H. performed the sequencing. R.B., K.R.H, A.V. and G.W.Y. analyzed data and wrote the manuscript.

## Acknowledgements

We would like to thank the members of the Ravits and Yeo labs for constructive discussions and comments on the manuscript. G.W.Y. was supported by grants from the National Institute of Health (HG004659, NS075449 and U54HG007005) and CIRM (RB1-01413 and RB3-05009). J.R. was supported by grants from the National Institutes of Health (NS051738), Microsoft Research, the Wyckoff family, the Moyer Foundation, Mrs Lois Caprile and the Benaroya Foundation. R.B. is a Myotonic Dystrophy Foundation (MDF) postdoctoral fellow. G.W.Y. is an Alfred P. Sloan Research Fellow.

## Accession Numbers

The NCBI GEO accession number for the RNA-seq and splicing sensitive microarrays data reported in this paper is **GSE76220**.

## Figure Legends

**Table 1. Sequencing statistics of control and sALS motor neuron samples.**

**Supplementary Figure 1.**
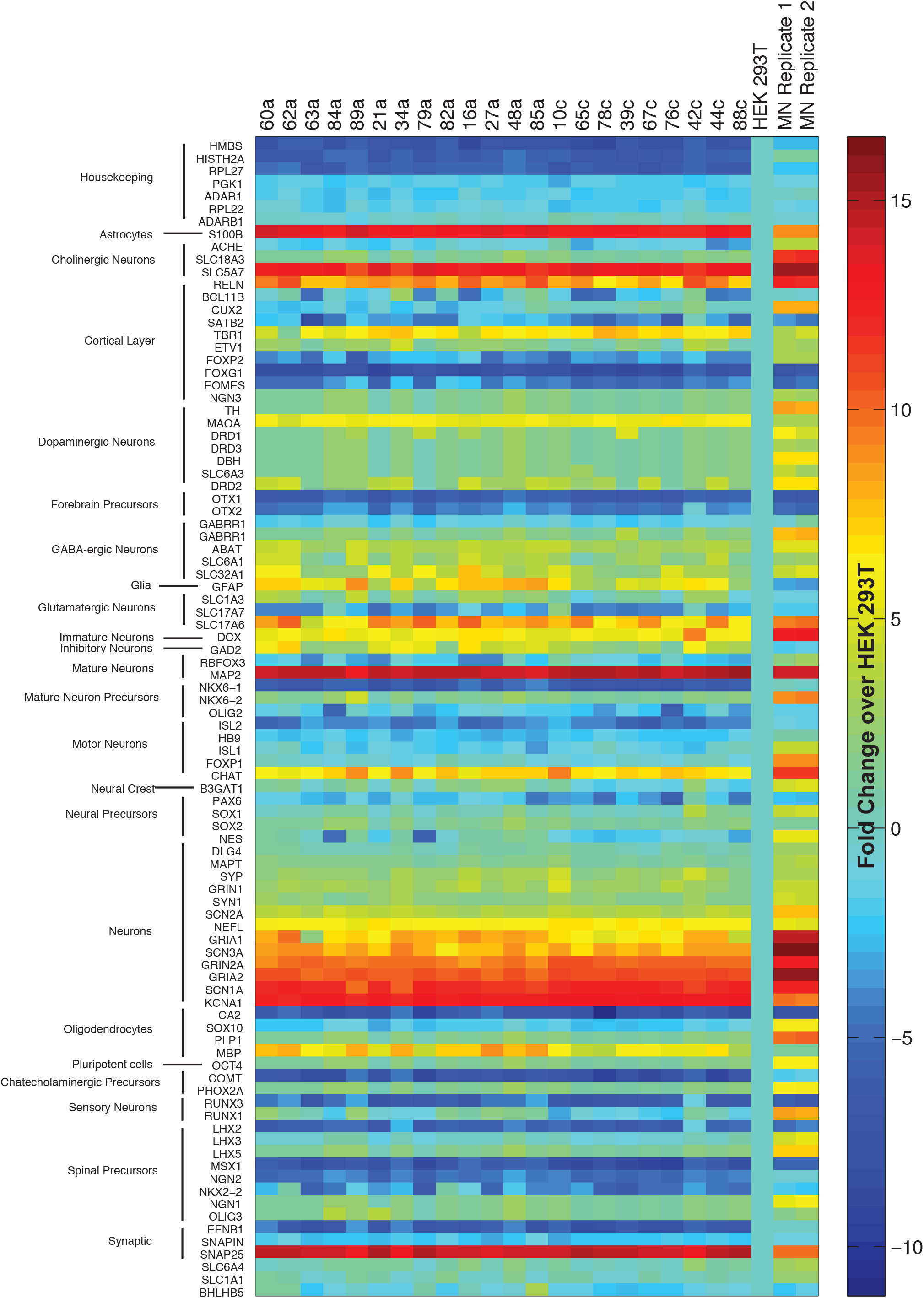
Motor neuron gene expression signature. Extended heat map of gene-expression in LCM microssected motor neurons normalized to HEK293T gene expression. Two iPSC derived motor neuron cell lines are also used as controls (right 2 lanes).

**Supplementary Figure 2.**
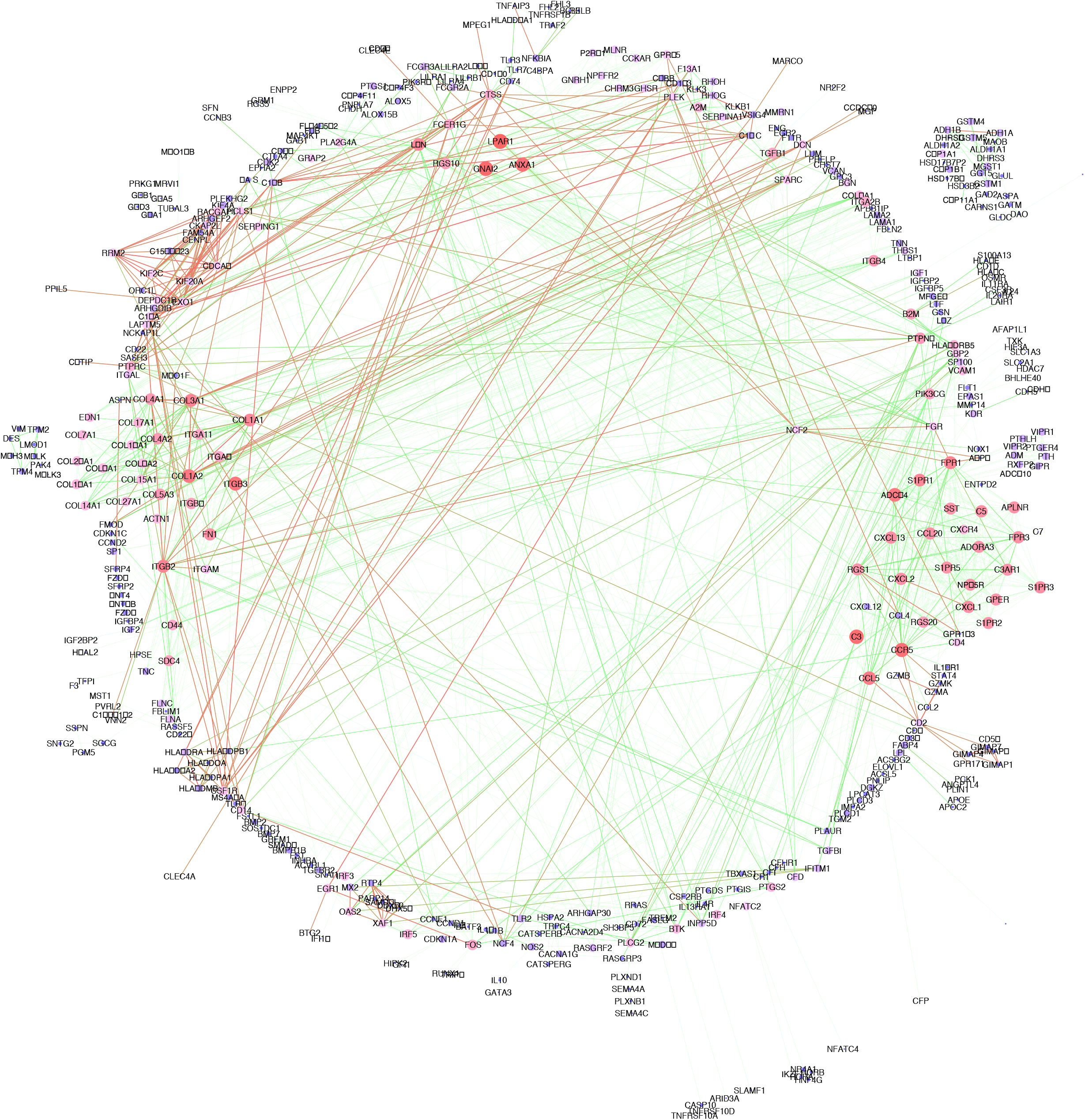
Gene interaction network analysis. Gene interaction network was constructed using MPR upregulated genes, STRING v10 tool, and Cytoscape. The nodes are colored according to degree or number of interactions for each node (red = high; dark blue = low). Red nodes are therefore hubs and are predicted to effect downstream pathways more. Edges are colored according to experimental co-expression (red = high; green = low). Red edges connect genes that are more likely to be co-expressed or co-affected.

**Supplementary Figure 3.**
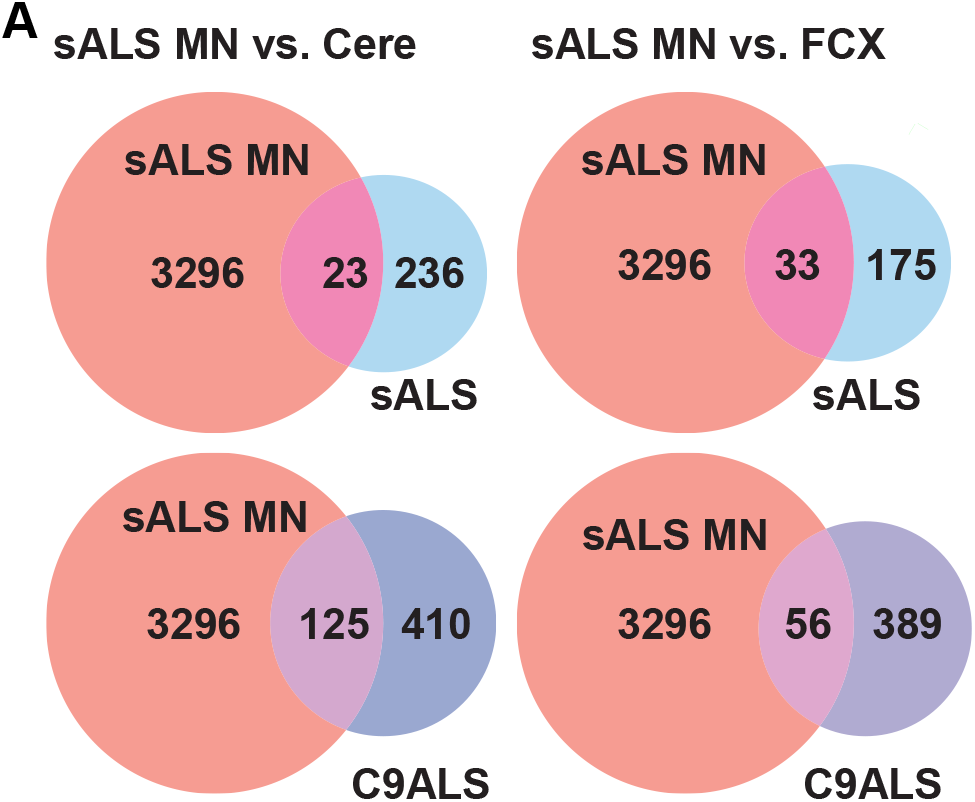
Intersection of sALS MN gene-expression data with existing ALS data. Venn diagrams showing overlap between sALS LCM MN gene-expression data and sALS and C9ALS data (frontal cortex and cerebellum) from Prudencio et al 2015 [9].

**Supplementary Figure 4.**
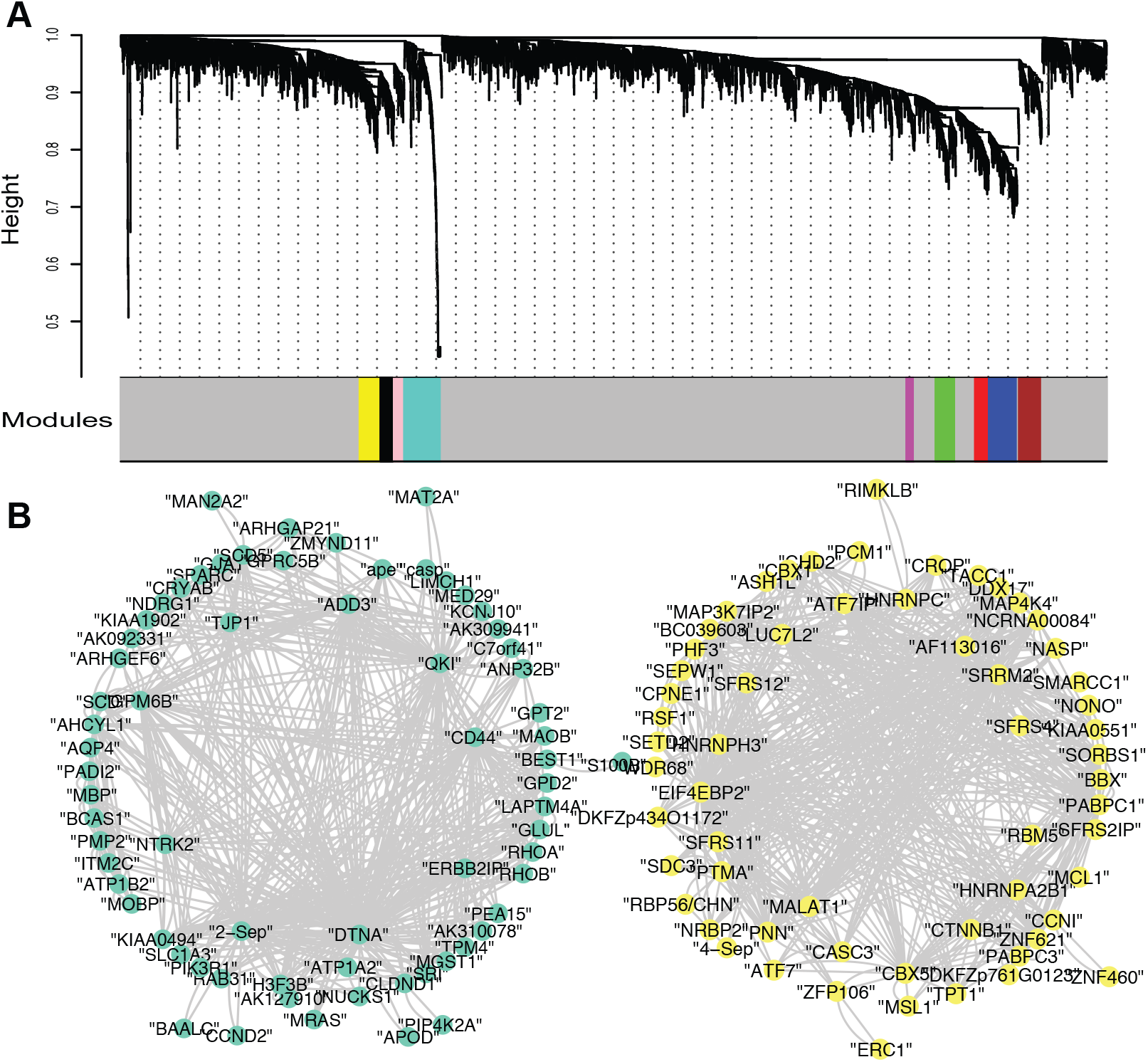
WGCNA co-expression network analysis. A) Dendrogram showing various modules detected by WGCNA analysis of sALS and control LCM MN data. (B) Network diagrams showing the genes in Turquoise and Yellow modules upregulated in sALS MNs. Hub genes (genes with greater connectivity) are depicted inside the larger circle of genes.

**Supplementary Figure 5.**
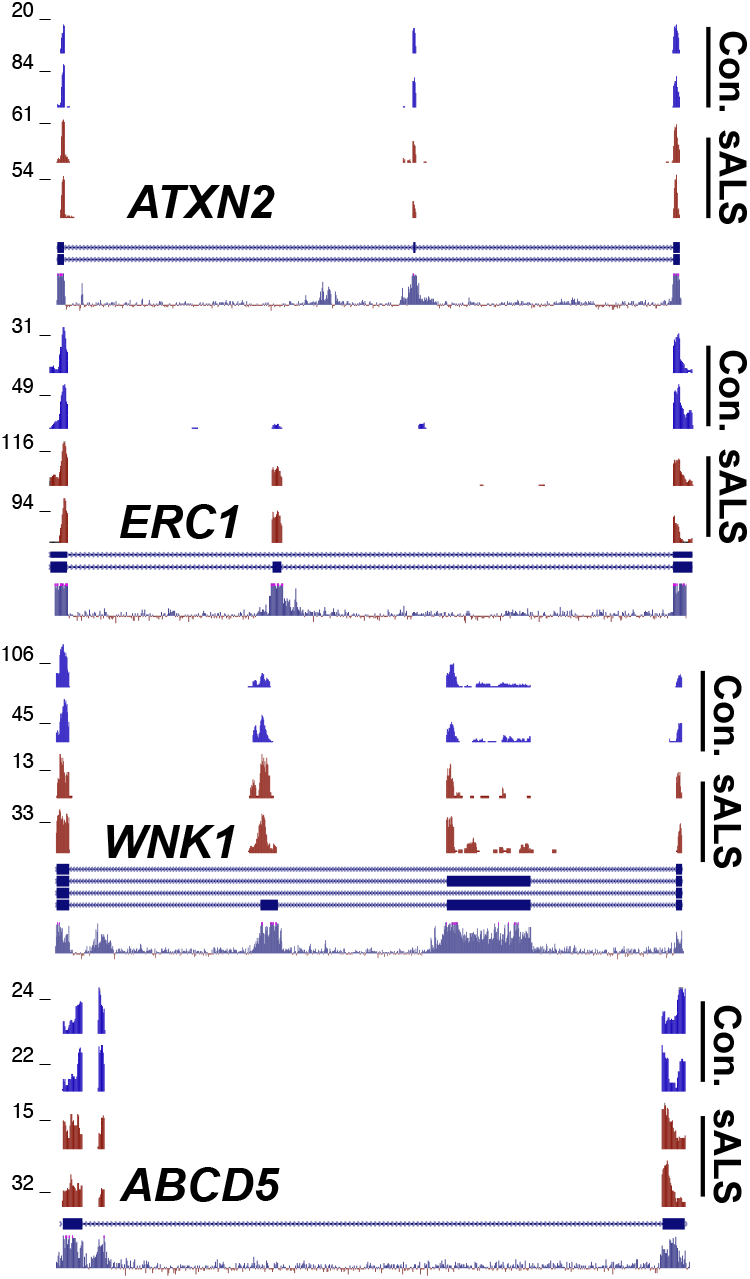
Splicing is misregulated in sALS motor neurons. Wiggle plots showing aberrantly regulated exons in sALS samples (red).

